# Bayesian and likelihood placement of fossils on phylogenies from quantitative morphometries

**DOI:** 10.1101/275446

**Authors:** Caroline Parins-Fukuchi

## Abstract

Jointly developing a comprehensive tree of life from living and fossil taxa has long been a fundamental goal in evolutionary biology. One major challenge has stemmed from difficulties in merging evidence from extant and extinct organisms. While these efforts have resulted in varying stages of synthesis, they have been hindered by their dependence on qualitative descriptions of morphology. Though rarely applied to phylogenetic inference, traditional and geometric morphometric data can improve these issues by generating more rigorous ways to quantify variation in morphological structures. They may also facilitate the rapid and objective aggregation of large morphological datasets. I describe a new Bayesian method that leverages quantitative trait data to reconstruct the positions of fossil taxa on fixed reference trees composed of extant taxa. Unlike most formulations of phylogenetic Brownian motion models, this method expresses branch lengths in units of morphological disparity, suggesting a new framework through which to construct Bayesian node calibration priors for molecular dating and explore comparative patterns in morphological disparity. I am hopeful that the approach described here will help to facilitate a deeper integration of neo- and paleontological data to move morphological phylogenetics further into the genomic era.

## Introduction

The role of fossil data in reconstructing phylogeny among living organisms has long been a central, yet contentious, topic in evolutionary biology. This has manifested over the past decade in the rapid proliferation of ’total-evidence’ methods that seek to simultaneously reconstruct the relationships and divergence times between living and fossil taxa using cladistic morphological matrices. These approaches, based upon probabilistic models of molecular and morphological character, have increased understanding of evolutionary tempo across large clades, and provide compelling evidence in favor of incorporating fossils in phylogenetic analyses (Pyron 2011; Ronquist *et al*. 2012). This can benefit both paleo- and neontological studies by improving the accuracy and treatment of uncertainty in estimation of divergence times and comparative dynamics (Slater *et al*. 2012; Guindon 2018).

A constant source of difficulty when jointly estimating phylogeny between living and extinct organisms is the unavailability of molecular data in nearly all fossil taxa. As a result, there has been a need to explore the compatibility of molecular with morphological data to better understand the capability of fossil and extant species to reciprocally inform reconstruction of phylogeny and divergence times. Previous work has sought to determine whether the inclusion of molecular data representing extant species can improve the reconstruction of relationships among fossils represented by morphology alone (Wiens 2009; Wiens *et al*. 2010). The results of these studies suggest that the inclusion of morphological characters comprising living and fossil species does not have a tendency to decrease the accuracy of phylogenetic reconstructions, and can improve estimation of fossil placements in well-behaved datasets. Expanding upon these observations, Berger and Stamatakis (2010) have shown that methods placing fossils on fixed molecular phylogenies can yield accurate results. Their study also shows that a scaffolding approach can further improve fossil reconstructions by offering a straightforward means of filtering through noise in morphological datasets by leveraging information from the molecular reference topology.

Morphological data present other unique challenges important to phylogenetic analysis. For example, morphological data are frequently susceptible to displaying biased or misleading signal. Although discordance in morphological datasets may sometimes reflect biological processes such as convergent evolution and hemiplasy, there is also frequently substantial noise stemming from systematic error and poor preservation of fossil taxa. Systematic sources of discordance often stem from the general practice of assigning discrete character states to taxa through qualitative assessment. The subjective nature of this process can cause major irreconcilable disagreement between results achieved from different researchers (Hauser and Presch 1991; Pleijel 1995; Wilkinson 1995; Hawkins *et al*. 1997; Scotland and Pennington 2000; Scotland *et al*. 2003; Brazeau 2011; Simões *et al*. 2017). As an added source of potential bias, these matrices are also frequently filtered to exclude characters that researchers suspect to be homoplasious. However, since these judgments are typically made subjectively, it may be of benefit to introduce a quantitative framework to evaluate the reliability of morphological traits.

As another challenge, the discrete character matrices most commonly employed in phylogenentics can often be difficult to adequately model. At present, researchers employing probabilistic methods generally use the so-called ‘Mk’ model (Lewis 2001). This is a generalization of the Jukes-Cantor model of nucleotide substitution that accommodates *k* possible character states. Although previous work based upon simulated data has suggested that Mk-based approaches outperform parsimony (Wright and Hillis 2014), the extent and conditions under which this is the case in empirical datasets is unclear (Goloboff *et al*. 2017). Empirical datasets are also likely to depart significantly from the assumptions of the Mk model. This poor match between model assumptions and data can lead to erratic results and high uncertainty in posterior estimates of divergence times (Ronquist *et al*. 2016). Although recent studies have proposed more sophisticated models (Wright *et al*. 2016), the standard symmetric Mk model remains in frequent use, and the sensitivity of topological reconstruction to this frequent mismatch is fairly unclear at present.

For all of these reasons, continuous traits have been suggested as a potential alternative (Felsenstein 1973, 1988; MacLeod 2002). Nevertheless, their use has remained relatively unexplored. In a previous study (Parins-Fukuchi 2017), I explored through simulations the relative performance of continuous and discrete traits in phylogenetic inference. I found that continuous characters perform similarly to discrete characters when phylogenetic half-life is set to be equal, while exploring the possibility that continuous traits may extend phylogenetic informativeness over some discretized character codings.

Traditional linear morphometric measurements have long been employed in morphological phylogenetics, but are typically discretized to more easily analyze them alongside present-absence data. Several approaches have been proposed for the discretization of quantitative morphological data (Thiele 1993; Wiens 2001). However, these can yield inconsistent or misleading results (Rae 1998; Goloboff *et al*. 2006), and may in principle reduce the amount of information in continuous datasets by binning fine-scaled variation into shared discrete categories. As a result, it may often be preferable to analyze continuous traits directly.

Tools that quantify morphological size and shape have the capacity to alleviate many of the concerns relating to bias and subjectivity that occur with discrete characters. Approaches such as geometric morphometrics offer the potential to holistically incorporate all dimensions of shape to inform phylogeny. The continuous state space of morphometric data might also increase the amount of information that can be extracted from morphological datasets, which may be beneficial when analyzing poorly-sampled fossil data. Continuous traits in general may engender benefits on two levels when available by 1) reducing subjective bias often encountered when constructing discrete character matrices, and 2) potentially preserving hard-won phylogenetic information over discretized character codings by representing the full range of observed interspecific variation. Although I explored point 2 previously (Parins-Fukuchi 2017), future studies will be needed to quantify the extent to which this is the case in diverse empirical datasets.

As another source of continuous traits, geometric morphometric data have shown utility in several previous phylogenetic studies using parsimony-based methods (González-José *et al*. 2008; Catalano *et al*. 2010; Smith and Hendricks 2013), but have not gained substantial traction. This may be in part due to the lack of available tools to analyze continuous trait data in a probabilistic framework. In addition, previous authors have raised concerns about the use of morphometric data in phylogenetic analysis, based primarily upon potential error stemming from covariance across characters and difficulties in parsing out homologous interspecific variation from variation resulting from rotations in morphospace (Felsenstein 2002). However, these concerns have been partially alleviated by the success of other workers in reconstructing phylogeny from landmark coordinates that are derived from truly homologous regions that have been properly aligned using Procrustes transposition (MacLeod 2001, 2002; Catalano *et al*. 2010; Goloboff and Catalano 2016).

The earliest studies investigating probabilistic methods of phylogenetic inference were developed using continuous characters modeled under Brownian motion (BM) (Cavalli-Sforza and Edwards 1967; Felsenstein 1973). Due in part to the abundant discrete character data that became available with the emergence of DNA sequencing, these approaches were quickly overshadowed in popularity by discrete trait approaches based upon Markov nucleotide substitution models. Continuous trait models have since gained significant popularity in phylogenetic comparative methods, but still are rarely used for phylogenetic inference. As a result, few implementations exist, with only ContML in the PHYLIP package and RevBayes providing such functionality (Höhna *et al*. 2016). However, the PHYLIP implementation uses a very simple tree searching procedure. RevBayes is very flexible, however, it is perhaps best suited to total-evidence analyses, where extant and fossil taxa are estimated simultaneously. An alternative procedure involves fixing extant relationships using the results of a molecular analysis, and estimating the positions of fossil taxa along this scaffolding. Previously, Revell *et al*. (2015) described a method that places individual taxa on phylogenies using quantitative data. The authors found that the approach performed well, but the implementation developed for the study was restricted to the placement of only extant and recently extinct taxa. In addition, the authors explored only the placement of a single taxon at a time.

Although, like the Mk model, BM is fairly simplistic, it may offer a degree of flexibility that improves its’ fit to empirical data in comparison to Mk. For instance, the Mk model assumes that stationary frequencies of character states are equal, whereas BM assumes that traits at the tip of a phylogeny are distributed according to a multivariate Gaussian distribution, with a set of covariances defined by the topology and branch lengths. While the Mk equilibrium assumption is violated in most empirical datasets, the BM assumption of normality can often be justified by the central limit theorem. This suggests that, even in cases where character state changes may better conform to a non-Gaussian distribution over short timescales, these collapse into a Gaussian-like distribution over longer timespans with many repeated draws. The standard phylogenetic BM model may still be violated by patterns such as directional change, but the effect is not well understood. Quantitative trait evolution might also proceed according to stasis and sudden jumps (Landis *et al*. 2013), but the identifiablility between BM and more complicated models across a tree when branch lengths are expressed in unit variance are not clear.

In this paper, I describe a new approach that places multiple fossils on molecular trees using quantitative characters modeled under BM. Departing from Revell *et al*. (2015), the phylogenetic BM model used here treats branch lengths in terms of morphological divergence rather than time. This simplifies the estimation procedure, and allows morphological disparity across taxa to be easily visualized across the resulting tree, similarly to molecular phylograms. The approach here seeks to tackle some of the most pressing obstacles associated with the use of traditional and geometric morphometric data in phylogenetic inference. Using simulated data, I validate and explore the behavior of the implementation. I also analyze empirical datasets representing the Vitaceae family of flowering plants (Chen 2009) and carnivoran mammals (Jones *et al*. 2015) comprised of traditional and geometric morphometric measurements, respectively. The method uses Markov chain Monte Carlo (MCMC) to infer the evolutionary placements of fossils and branch lengths.

## Methods and Materials

### Software

All fossil placement analyses were performed using the new software package *cophymaru* written in the Go language. The source code is publicly available as free software at https://github.com/carolinetomo/cophymaru. This package estimates the positions of fossil taxa on a user-specified reference tree of extant species using continuous traits contained within a PHYLIP-formatted data file where each trait is separated by tabs. Examples can be gleaned from the simulated and empirical data generated from this study, available online.

### Brownian motion model

The approaches that I describe in this paper all rely upon the familiar BM model of evolution (Butler and King 2004; O’Meara *et al*. 2006). Under BM, traits are assumed to be multivariate distributed, with variances between taxa defined by the product of their evolutionary distance measured in absolute time and the instantaneous rate parameter (*σ*):

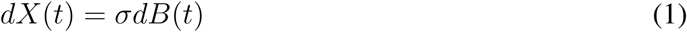

where *dX(t)* is the time derivative of the change in trait *X* and *dB(t)* corresponding to normally distributed random variables with mean 0 and variance *dt*. This leads to the expectation that over time *t*,

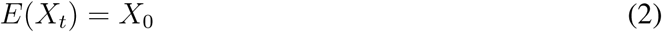

with

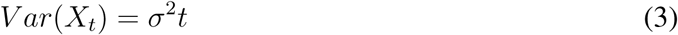

where *X*_0_ gives the trait value at *t*_0_.

The methods that I describe use a slightly different parameterization and likelihood calculation than most conventional implementations used in modern phylogenetic comparative methods (PCMs). These generally construct a variance-covariance (VCV) matrix from a dated, ultrametric phylogeny to calculate the likelihood of the data, assuming a multivariate normal distribution (Butler and King 2004; O’Meara *et al*. 2006). Since these methods treat the topology and branching times as known, the goal is typically to obtain the maximum likelihood estimate (MLE) of the rate parameter *(σ*) to examine evolutionary rate across clades.

In typical usage, researchers employ phylogenetic BM models where branch lengths are scaled to absolute time, and a rate parameter is estimated. Although it is possible to simultaneously estimate divergence times and topology while analyzing continuous traits, this requires the specification of a tree prior that can accommodate non-ultrametric trees that include fossils. In addition, this approach would effectively perform morphological dating using continuous traits. The behavior and feasibility of such a procedure is not understood, and falls outside the scope of this article. Perhaps more importantly, this would also create circularity when using the method to place fossils used as calibrations in molecular dating. To overcome the need for simultaneously estimating divergence times and fossil placements, the method estimates the product *σ*^2^*t* together. As a result, rate and absolute time are confounded in the trait and tree models. Branch lengths, which reflect the morphological disparity between taxa, are thus measured in units of morphological standard deviations per site. This interpretation could be thought roughly of as a continuous analogue to the branch lengths obtained from discrete substitution models. Similarly to the discrete case, long branch lengths could reflect either a rapid rate of evolution or a long period of divergence (in absolute time) along that lineage.

### Computation of the likelihood

Rather than use the computationally expensive VCV likelihood calculation, I use the reduced maximum likelihood (REML) calculation described by Felsenstein (1973). Full derivations of the likelihood and algorithm are also given by Felsenstein (1981b) and Freckleton (2012), and summarized briefly below. The tree likelihood is computed from the phylogenetic independent contrasts (PICs) using a ‘pruning’ algorithm. In this procedure, each internal node is visited in a postorder traversal, and the log-likelihood, *L_node_* is calculated as multivariate normal, with a mean equal to the contrast between the character states, *x*_1_ and *x*_2_ at each subtending edge and variance calculated as the sum of each child edge, *v*_1_ and *v*_2_:

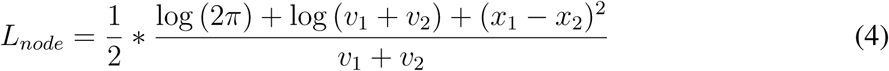

The PIC, *x_internal_*, is calculated at each internal node and used as the character state representing the internal node during the likelihood computation at the parent node. The edge length of the internal node, *v_internal_* is also extended by averaging the lengths of the child nodes to allow the variance from the tips to propagate from the tips to the root:

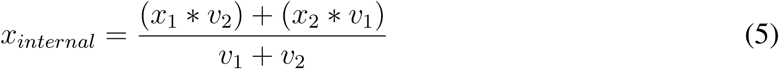

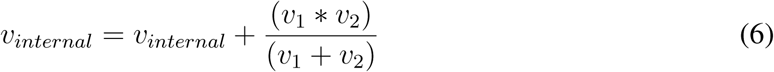

The total log-likelihood of the tree, *L_tree_* is calculated by summing the log-likelihoods calculated at each of the *n* internal nodes.

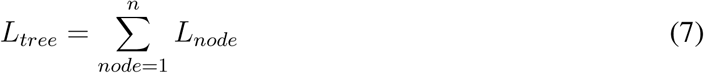

### Priors

Since the estimation of branch lengths from continuous traits is relatively uncharted territory in phylogenetics, I implemented and tested three different branch length priors derived from the molecular canon: 1) flat (uniform), 2) exponential, and 3) a compound Dirichlet prior after (Rannala *et al*. 2011). The compound Dirichlet prior also offers the option to set the scale of the expected tree length using the initial rough estimate of branch lengths.

### Markov-chain Monte Carlo

This method uses a Metropolis-Hastings (MH) algorithm (Hastings 1970) to simulate the posterior distribution of fossil insertion points and branch lengths. Rearrangements of the topological positions of fossil taxa are performed by randomly pruning and reinserting a fossil taxon to generate a proposal. This is a specific case of the standard subtree pruning and regrafting (SPR) move for unrooted tees (Fig. 1). In this procedure, the two edge lengths that link the fossil to the rest of the tree are merged when the fossil tip is pruned, while the edge upon which the tip is inserted is split into two. The move is described in detail, along with a full derivation of the appropriate MH proposal ratio in (Yang 2014, p. 287). Branch lengths are updated both individually and by randomly applying a multiplier to subclades of the tree. MH proposal ratios for branch length updates follow the derivations given for the the ’multiplier’ or ’proportional scaling’ move described by (Yang 2014, p. 225).

**Figure 1.**
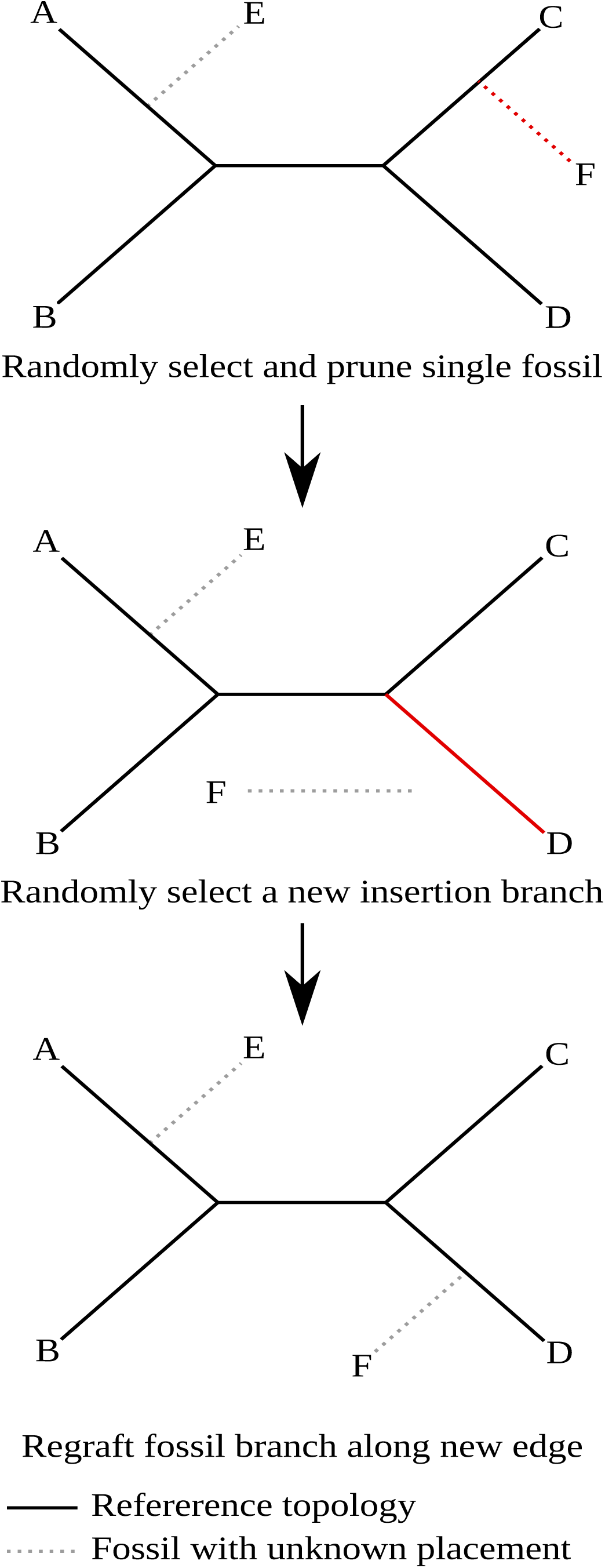
Random fossil prune and regraft procedure.

### Generating a rough ML starting tree

I re-implemented the approach used in the ContML program to generate an approximate ML starting tree. These initial placements are achieved using stepwise addition. Unlike ContML, this step successively adds fossils to the molecular guide tree, and so only the fossil positions are estimated. Each fossil is individually inserted along all existing branches of the tree, with the insertion point that yields the highest likelihood retained. At each step, MLEs of the branch lengths are computed using the iterative procedure introduced by (Felsenstein 1981a). In this procedure, the tree is rerooted along each node. PICs are calculated to each of the three edges subtending the new root, and are treated as ’traits’ at the tips of a three-taxon tree. The MLE of each edge length of the pruned three-taxon tree (*v*_i_) is computed analytically using the expressions::

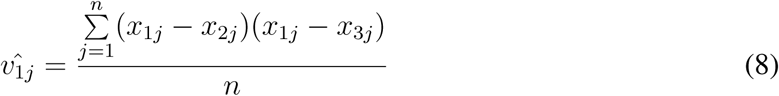

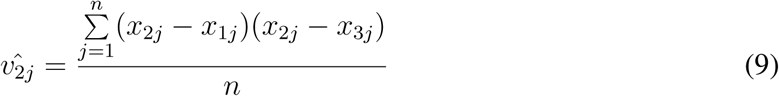

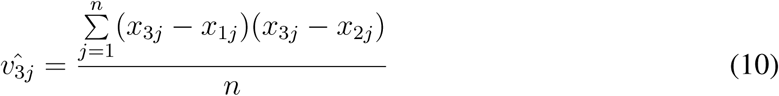

This process is iterated by successively rerooting on each node of the tree and calculating the branch lengths until their values and the likelihoods converge. Felsenstein (1981) gives a more detailed explanation of the algorithm, along with a complete derivation of the MLE branch length calculations.

Once an initial placement has been assigned for all of the fossils, the branch lengths are optimized on the complete tree. These starting lengths can be used to inform branch length priors used during MCMC simulation. One problem with interpreting the results of the ML approach on their own is that it has a strong propensity to becoming trapped in local optima. As a result, it should interpreted cautiously, and not used without further MCMC searching. In the applications here, the topologies achieved from this procedure are used only to construct starting trees, while the branch lengths inform the specification of branch length priors. This procedure allows straightforward construction of non-random starting trees for the MCMC and priors that reflect the the dataset under analysis.

### Filtering for concordant sites

One major hurdle involved in the use of morphological data is their frequent tendency to display noisy and discordant signal. This problem might be expected to manifest even more intrusively in morphometric datasets than in discrete datasets, since traits are much less likely to be excluded *a priori* on the basis of perceived unreliability. As a result, there is a need to filter through noisy signal to favor more reliable sites. I developed a procedure adapted from Berger and Stamatakis (2010) for this purpose. This computes a set of weights based upon the concordance of each site with the reference tree. In this procedure, the likelihood (*L*_ref_) of each site is calculated on the reference tree (excluding fossil taxa). Next, the likelihood (Ln) of each site is calculated along each *n* of 100 phylogenies generated randomly by successively grafting nodes in a stepwise manner until a full tree is formed. Branch lengths are then assigned using uniform random draws. If the likelihood of the site is higher along the reference tree than the current random tree, the weight of the site is incremented by one. Thus, site *j* receives the integer weight:

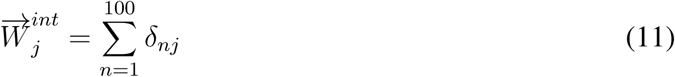

where *δnj* = 1 if:

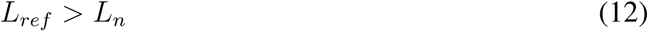

and *δnj* = 0 if:

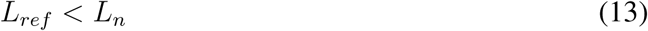

This yields a weight vector that is the same length as the character matrix, with each site possessing a weight between 0 and 100. The sites are then weighted using one of three schemes: 1) whole integer values, where the weight equals the value obtained from equation 7, 2) a floating point value between 0 and 1, where the value generated from the random comparison is divided by 100, and 3) a binary value where the weight is equal to 1 if the site displayed a higher likelihood in the reference tree than 95 or more of the random trees, and 0 if less than 95:

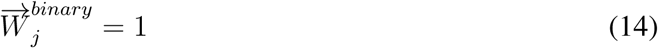

if

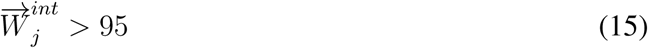

and

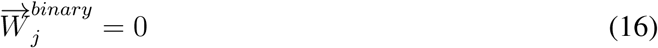

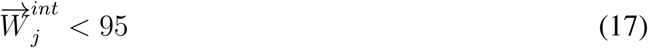

After the weights are computed using the input guide tree, they are stored, and used in all subsequent likelihood computations during MCMC simulations.

In application, I found that integer weighting caused poor MCMC mixing, and so the floating and binary schemes are probably most practical in most cases. The poor mixing achieved by the integer scheme is likely due to the large increase in the scale of the log-likelihoods. This causes nearly all proposals to be rejected, substantially reducing the efficiency of the algorithm. In effect, the MCMC algorithm becomes a very inefficient hill-climbing ML search, since only proposals that increase the likelihood are accepted. Since it filters out discordant sites completely, the binary scheme is enforces a harsher penalty than the floating and integer schemes, and so might be of greatest use in particularly noisy datasets. As an additional note, although these procedures share similar terminology to the site weights calculated during parsimony analysis of multi-state characters, they differ in their purpose. Parsimony site weights are intended to normalize the contribution of characters with differing state spaces to the overall tree length. In contrast, the site weighting approach deployed here is designed to decrease the contribution of sites that disagree with the reference topology to the overall tree likelihood, instead highlighting signal taken to be more reliable. As a result, the guide tree is used to identify sites that are most likely to reliably inform fossil placements.

Although this procedure was originally implemented in an ML context, the application here functions as a prior. By assuming that the molecular guide tree provides an accurate view of extant species relationships, characters that appear to show significant error, homoplasy, or reflect other processes yielding discordant signal, are filtered out or de-emphasized. This procedure has the effect of increasing posterior support in datasets possessing many discordant characters. The Bayesian framework offers a straightforward means to interpret the resulting support values as standard posterior credibility estimates. Nevertheless, the filtering approach, as any prior, should be applied thoughtfully, and compared to results when the prior is not used.

### Simulations

To explore the behavior of these approaches under different settings and validate the implementation, I performed a set of simulations. From a single simulated tree, I pruned five “fossil” taxa and estimated their positions along the tree using 100 datasets of 50 characters simulated under BM. The tree was simulated under a birth-death model, with a birth parameter of 1.0 and a death parameter of 0.5. The resulting tree conained 41 taxa, leaving a 36-taxon reference tree when the five fossils were pruned. To explore the effect of conflicting and noisy signal, I also generated alignments consisting of 50 “clean” traits simulated along the true tree, and combined with sets “dirty” traits in intervals of 10, 25, and 50 traits generated along random trees. All trait (clean and dirty) simulations were performed using the “fastBM” function in the phytools package (Revell 2012). All traits were simulated using a rate parameter of 1.0. Random trees were generated by collapsing the true tree into a star topology using the “di2multi” function, which was randomly resolved using the “multi2di” function. Branch lengths were than assigned randomly by drawing from an exponential distribution with mean set to 1. The simulated data sets and all scripts used to generate them are available at https://github.com/carolinetomo/fossil_placement_tests.

I restricted the simulations to a fairly small number of traits because this reflected a similar size as the two empirical datasets. This level of sampling is fairly common among existing continuous datasets, which are typically compiled from only one or two organs. In the future, I am hopeful that quantitative morphometric datasets will increase in size to encompass much broader sampling across organs, but at present, it seemed most sensible to examine the level of sampling expected from existing datasets. Unlike in a previous paper (Parins-Fukuchi 2017), I also did not simulate traits that display covariance among sites. This is because 1) I showed in the previous study that covariance does not in and of itself significantly handicap reconstructions from continuous traits, and 2) because in this study I was primarily interested in examining the effect of inducing random noise without the potentially confounding effect of covariance. Although covariance has been expressed as a major concern in morphometric phylogenetics (Felsenstein 1988, 2002), there is no reason to expect greater covariance between continuous traits than discrete traits, which, ideally, should describe similar aspects of morphology. Nevertheless, a fairly common source of error in molecular phylogenetic studies can occur when many sites exhibit shared misleading signal due to some legitimate biological process. A similar effect may in principle occur in studies using continuous morphological characters. And so, although continuous trait matrices may not necessarily carry greater inherent risk toward being mislead by covariance across sites than studies based on molecular and discrete morphological characters, careful analysis is important to properly dissect the distribution of signal across character matrices to properly identify biological and systematic sources of conflict and error.

These simulated datasets were then used to reconstruct the placements of the five fossils. To explore the relative performance of weighting schemes, I performed reconstructions using both the binary and floating approaches. These were supplemented by analyses of the noisy datasets without applying site weights. MCMC simulations were run for 1,000,000 generations and checked to ensure that the effective sample sizes (ESS) exceeded 200. The exponential branch length prior was employed for the simulated data with a mean of 1.0. To evaluate the accuracy of the placement method, I then calculated the distances between the true and reconstructed fossil placements. This was calculated by counting the number of nodes separating the true insertion branch from the reconstructed insertion branch. Placement accuracy was evaluated using the *maximum a posteriori* (MAP) summaries of tree distributions. MAP trees represent the single most sampled tree during the MCMC run. Tree summary and placement distances were calculated using custom Python scripts.

### Empirical analyses

To assess the utility of the new approach in analyzing continuous morphological data, I performed analyses on empirical datasets comprised of 1) linear measurements and proportions, and 2) geometric morphometric data composed of 3-dimensional landmark coordinates. These are two common sources of continuous trait data, and so were chosen to test the method across different possible data types. In the *cophymaru* implementation of the method, these characters are input as character matrices similar to those used to store discrete traits, with homologous measurements arranged in columns, corresponding to rows of taxa. In the case of the geometric morphometric data, each landmark coordinate represents a column, similarly to previous phylogenetic approaches that explicitly use geometric morphometric data (Catalano *et al*. 2010). Empirical character matrices, trace files, and reference trees are all available online at https://github.com/carolinetomo/fossil_placement_tests.

I estimated the phylogenetic positions of fossils using a morphological matrix comprised of 51 continuous measurements gathered from pollen and seed specimens sampled across 147 extant and 8 fossil Vitaceae taxa. These data were acquired from Chen (2009). I constructed a guide tree for the extant taxa from 8 nuclear and chloroplast genes gathered from Genbank using the PHLAWD system (Soltis *et al*. 2011). The sequence alignment used to construct the guide tree is available in the online data supplement. Using this scaffolding, I analyzed the morphological data to estimate the positions of the fossil taxa. Individual runs were performed under all three branch length priors to assess stability across models. All analyses were run for 30,000,000 generations and visually checked for convergence. Analyses were performed with binary weights applied to the sites and compared to an unweighted analysis. To ensure that MCMC runs were not trapped in local optima, several redundant runs were performed under each combination of settings. For each, the analysis with the highest mean likelihood was retained.

To explicitly test the informativeness of geometric morphometric data in fossil placement, I also performed analyses on a dataset of 33 3D landmark coordinates representing 46 extant and 5 extinct fossil carnivoran crania (Jones *et al*. 2015). A reference tree composed of the 46 extant taxa was obtained from the data supplement of the original study. These coordinates were subjected to Procrustes transposition using MorphoJ (Klingenberg 2011). This yielded a matrix where each character represented the aligned X, Y, or Z position of one landmark. These characters are ’aligned’ such that each column contains the coordinates in one dimension of a single landmark occupied by each taxon. Although the details surround the analytical approaches differ, this use of morphometric data is similar to that used in the method described by Catalano *et al*. (2010). The resulting traits displayed phylogenetic signal, but the transposed coordinates showed very low dispersion (variance) on an absolute scale. Low variance can result in narrower peaks in the MCMC surface, which causes difficulties in achieving MCMC convergence. To remedy this, I scaled all of the traits to increase the absolute variance evenly across taxa evenly at each site while maintaining the original pattern of relative variances across taxa using the scale() function in R (R Core Team 2016). This procedure preserved the signal present in the original dataset, since the relative distances between taxa remained the same. Final analyses were performed on this transformed set of measurements. As with the Vitaceae dataset, I analyzed the canid data under all three branch length priors, and performed several runs, retaining the one with the highest mean likelihood. MCMC simulations were run for 20,000,000 generations, and visually examined using Tracer v1.6 to assess convergence. Both empirical datasets achieved large ESS values (>1000) under all settings.

For both datasets, I used starting trees and branch lengths generated from the rough ML method described above. Sites were weighted using the binary for the final analyses. Intermediate analyses using unweighted and float-weighted sites were also performed, and are presented in the data supplement. Dirichlet priors were assigned alpha parameters of 1.0 and beta parameters specified as the total tree length of the ML starting tree. Exponential branch length priors were assigned mean values of 1.0.

Since the empirical datasets were more complex than the simulated data, I summarized the tree distributions as maximum clade credibility (MCC) summaries. These summaries maximize the support of each clade. These were compared to the MAP estimates, however, and yielded generally concordant placements (supplementary material). MCC summaries were obtained using the SumTrees script that is bundled with the DendroPy package (Sukumaran and Holder 2010). Branch lengths were summarized as the mean across all sampled trees.

## Results and Discussion

### Simulations

Reconstructions of fossil placements from the simulated datasets showed that the method is generally accurate in placing fossil taxa (Tables 1 and 2). In the absence of noisy traits, reconstruction is nearly always correct, with the reconstructed position of each fossil placed less than 0.1 nodes away from the true position on average. In the presence of random noise, the reconstructions are fairly accurate, except when noise becomes severe. Nevertheless, even in the extreme case where half of the characters display completely random signal, the estimated fossil positions tend to fall within the correct region of the tree, falling 1.85 and ~3 nodes away from the correct placement on average under the exponential prior when alignments contain an equal number of clean and dirty sites. And although the procedure reconstructs fossil positions that are quite distant in the worst case (3 nodes away, on average), application of the weighting procedures improves reconstructions, even though the signal-to-noise ratio is quite high.

**Table 1.** Mean distances of true and reconstructed fossil placements under the exponential branch length prior. Distances are measured as the average number of nodes separating reconstructed placements from their true positions across all 100 replicates of each dataset.

| dataset | binary_weights | float_weights | unweighted |
| --- | --- | --- | --- |
| 50 clean | 0.065 | 0.067 | 0.059 |
| 50 clean + 10 dirty | 0.156 | 1.234 | 2.516 |
| 50 clean + 25 dirty | 0.965 | 2.623 | 3.105 |
| 50 clean + 50 dirty | 1.85 | 3.069 | 3.348 |

**Table 2.** Mean distances of true and reconstructed fossil placements under the compound Dirichlet branch length prior. Distances are measured as the average number of nodes separating reconstructed placements from their true positions across all 100 replicates of each dataset.

| dataset | binary_weights | float_weights | unweighted |
| --- | --- | --- | --- |
| 50 clean | 0.054 | 0.053 | 0.072 |
| 50 clean + 10 dirty | 0.131 | 1.192 | 2.466 |
| 50 clean + 25 dirty | 0.751 | 2.552 | 3.175 |
| 50 clean + 50 dirty | 1.474 | 2.982 | 3.324 |

In the *cophymaru* implementation, the compound Dirichlet prior outperforms the exponential branch length prior on the simulated datasets (Table 2). The mean distance between the true positions of fossils and their inferred placements is smaller under the compound Dirichlet in all but one of the comparisions. The improvement exhibited under the compound Dirichlet is greatest when using the binary weighting scheme, resulting in placements approximately 0.2–0.4 nodes closer to the true position across all simulated datasets. The improvement also increases with the noisiness of the simulated dataset, with the 50 clean+50 dirty dataset displaying the largest increase in placement accuracy. This result suggests that the compound Dirichlet branch prior combined with binary weighting scheme may be the ideal mode through which to analyze particularly noisy datasets.

Across both branch length priors, binary weighting shows improved accuracy over float and unweighted analyses. However, despite the apparent advantage of binary weighting, it is possible that the float weighting scheme could remain beneficial in cases where the distribution of noise varies betwen different regions of trees. This is because the float weighting scheme limits the contribution of noisy sites to the likelihood rather than entirely excluding them. This possibility was not examined in this set of simulations, since the dirty traits were generated to reflect completely random noise. However, in reality, noise may be structured to display discordance in only certain taxa. In these cases, continuous traits may display misleading signal among some subset of taxa, but correctly informative signal among other subsets. Further work will be needed to determine the extent to which weights calculated under the float weighting scheme vary when conflict is localized to particular regions of the reference tree.

Overall, the simulations demonstrate the efficacy of the method for the phylogenetic placement of fossils and provide a validation of the computational implementation. The analysis of clean datasets shows that the method performs well, estimating fossil placements with very low error when signal is clear. The adaptation of Berger and Stamatakis’ (2010) site weight calibration approach also appears to effectively filter through noisy datasets to improve estimation. The binary weight calibrations appear particularly effective at dealing with rampant misleading random noise, with improving accuracy by 2 to 20 times depending on the relative proportion of signal and noise compared to unweighted analyses. These results show promise toward the prospect of applying the method developed in this work to the analysis of large-scale morphometric datasets, where significant noise might be expected. Although introducing noise to unweighted analyses decreases reconstruction accuracy, the method performs predictably, and still manages to place fossils on average within the correct neighborhood. However, when weighting schemes are applied, the performance improves drastically, highlighting the promise of this method for the analysis of empirical datasets.

### Vitaceae dataset

Application of the fossil placement method to the Vitaceae dataset showed generally positive results (Fig. 2, Fig. S1). The weight calibration procedure revealed substantial noise in the dataset, with 10–12 of 51 sites failing to favor the molecular reference tree over the random trees at least 95% of the time across all runs. Despite this noise, the binary weighting scheme appeared to adequately filter through this noise to generate biologically reasonable results. *Vitis tiffneyi, Parthenocissus_clarnensis*, and *Ampelopsis rooseae* all share clades with the extant members of their respective genera. *Palaeovitis_paradoxa*, and *Cissocarpus jackesiae*, which represent genera with no extant species, both group with separate, non-monophyletic groups of crown *Cissus. Ampelocissus wildei* placed within crown *Cissus*, separated by only a node from *Palaeovitis paradoxa*. All six of these taxa are stable in their placements, grouping within the same clades across runs, and when both the exponential and empirical compound Dirichlet priors are applied.

**Figure 2.**
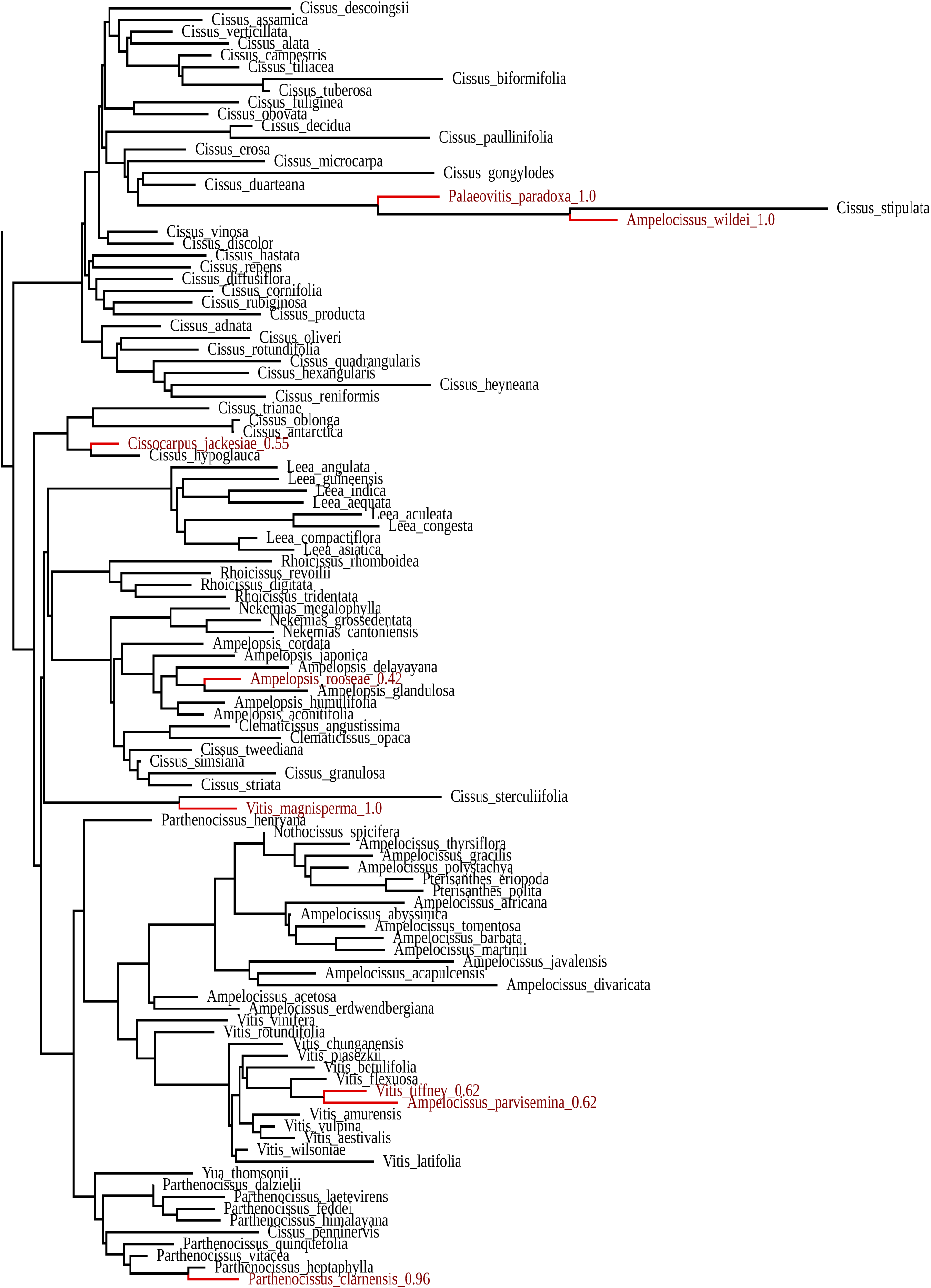
Vitaceae fossil placements inferred under the compound Dirichlet branch length prior. Fossil taxa and branches are highlighted in red. Values following fossil tip labels indicate posterior support for placement. Topology is summarized from the posterior using the set of maximally credible clades (MCC). Figure displays only the clade containing all 6 fossils. The full Newick tree is available in the data supplement.

The remaining two fossils are unstable in their placements across branch length priors. *Ampelocissus parvisemina* alternately occupies clades shared by crown *Vitis* or *Nekemias* in the exponential and Dirichlet prior runs, respectively. This taxon shows poor support under the exponential prior, and achieves higher posterior support under the compound Dirichlet prior. Under the exponential prior, the *Ampelocissus parvisemina* placement shows a 0.2 posterior probability (Fig. S1), and increases to 0.62 under the Dirichlet prior (Fig. 2). Similarly, *Vitis magnisperma* alternately resolves into clades shared by crown *Cissus* and *Ampelocissus* under the exponential and Dirichlet priors, with posterior support values of 0.23 and 0.54, respectively.

The simulations show that the compound Dirichlet prior achieves higher accuracy than the exponential prior, especially when combined with the binary scheme and applied to noisy datasets. If this observation can be extended to the empirical results, it is reasonable to prefer the placements inferred for these two taxa under the compound Dirichlet prior. This interpretation is supported by the greater stability and higher posterior support observed under the compound Dirichlet branch length prior.

### Carnivoran dataset

Analysis of the carnivoran dataset also yielded generally reasonable results (Fig. 3). The placements of *Piscophoca pacifica, Acrophoca longirostris, Enaliarctos emlongii*, and *Allodesmus* agree with previous results (Amson and de Muizon 2014; Jones *et al*. 2015). The placement of *Piscophoca pacifica* and *Acrophoca longirostris* differs slightly from the topology generated by Jones *et al*., placing the two taxa in a more nested position. However, this placement is consistent with the results of Amson and Muison. *Enaliarctos emlongii* and *Allodesmus* resolve in positions identical to the topology used by Jones and colleagues (2015). *Pontolis magnus* is more erratic in its placement, alternating between placement at the center of the unrooted topology, or grouping with *Vulpes* and *Otocyon*. The latter placement is unlikely to be correct, because it places *Pontolis magnus* within the Canidae family, while is canonically known as the only extant member of family Odobenidae. Nevertheless, like the problem taxa in the Vitaceae example above, the placement of *Pontolis* displays reassuringly weak support, both in terms of its posterior density and in its tendency to group at the center of the tree. Interestingly, although the placements of *Enaliarctos emlongii* and *Allodesmus* remain stable across runs, both display weak support.

**Figure 3.**
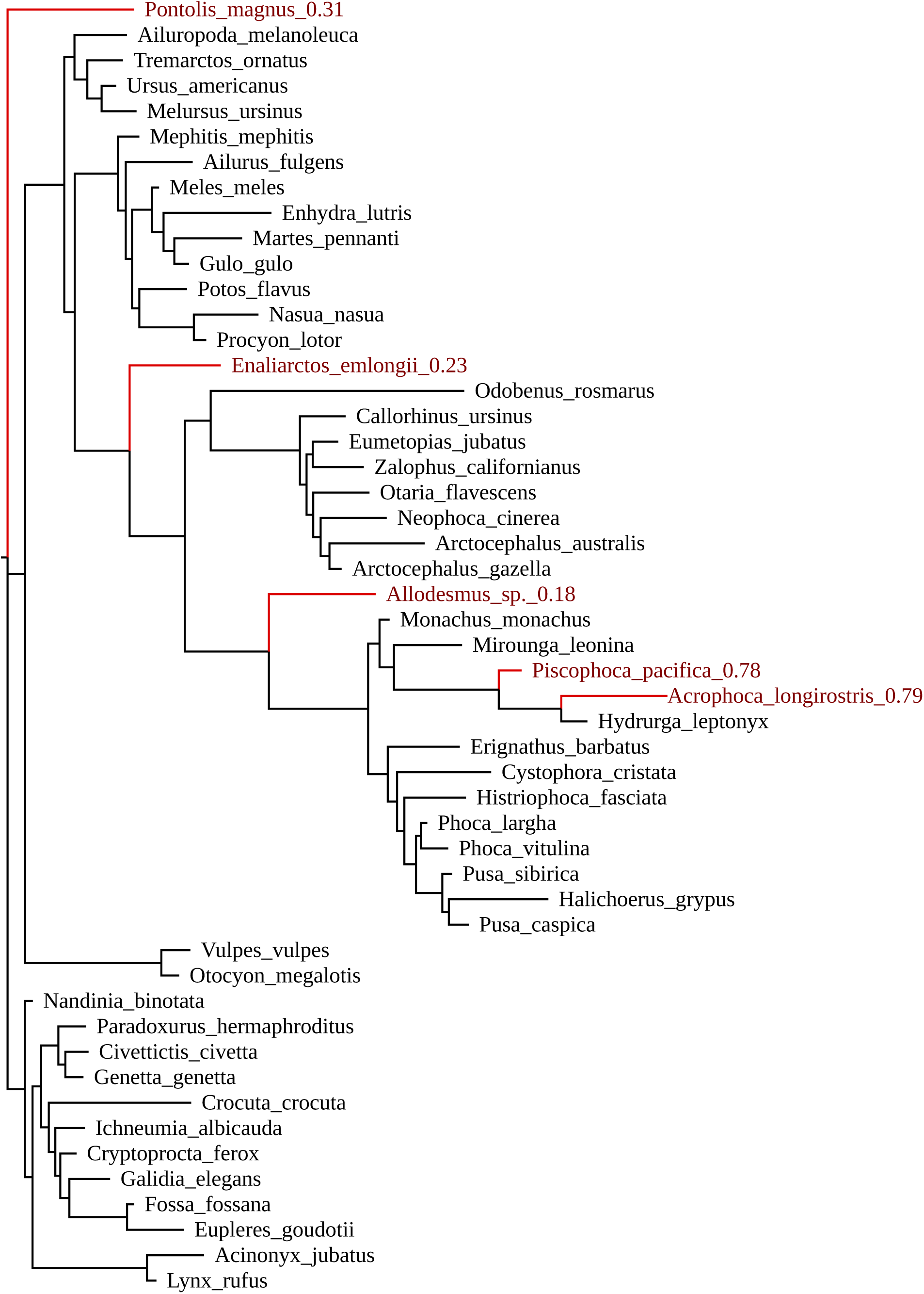
Fossil placements inferred from the carnivoran dataset using the compound Dirichlet prior. Placements are displayed as the maximum clade credibility summary of the posterior distribution of trees. Branch lengths represent morphological disparity. Values trailing fossil tip names display posterior support.

In both datasets, placement under the exponential branch length prior yields conservative estimates of uncertainty in the fossil placements, displaying generally low posterior support, except when placements are exceptionally stable such as with *Ampelocissus wildei*. This is especially important in ‘rogue’ taxa such as *Vitis magnisperma*. Branch support under the compound Dirichlet prior is higher across several fossils in the Vitaceae dataset. The positions of the six taxa with stable behavior (listed above) do not change significantly under the compound Dirichlet compared to the exponential prior. Closer examination is needed to better determine the significance of this apparent sensitivity of posterior support measures to prior choice observed in Vitaceae. The carnivoran dataset does not exhibit the same behavior, with both branch support and fossil placements similar across priors.

### Continuous vs discrete morphological characters

Previous work investigating the degradation of phylogenetic signal over time has implied that continuous traits can benefit over discrete traits under certain circumstances (Revell *et al*. 2008). In principle, methods that analyze continuous traits directly are preferable over those that bin continuous variation into discrete categories (Goloboff *et al*. 2006), due to their avoidance of error stemming from discretization schemes (Rae 1998), and potential to better preserve information that can be gleaned from morphological datasets (Parins-Fukuchi 2017). Nevertheless, depending on the type of continuous data that are used, the incorporation of features that can be uniquely described qualitatively, such as the loss and gain of structures, may be helpful. It would be straightforward to combine such discrete information into the morphometric framework described here. As progress in this area develops, it will be important to better understand the behavior of different sources of morphological data at different timescales, and the most appropriate ways to combine, model, and gather such datasets.

### Caveats to the approach

Although the performance of this new approach on simulated and empirical data appears generally promising, there are several caveats to consider in its use. When applying this method to geometric morphometric data, authors should be cautious to properly align landmark coordinates using Procrustes transformation to remove the effects of rotation in 3D space as a source of variation. In addition, as is shown by the simulations, when the signal-to-noise ratio becomes high, the weighting procedure performs significantly less accurately than when the amount of noisy/misleading signal is lower. Further work will be needed to assess the source of this discrepancy, and the possibility of additional steps that fortifies the approach when noise becomes high. The weighting procedure also may become more complicated in cases where a reliable scaffolding tree cannot be estimated due to genealogical discordance, or where signal displayed by the quantitative traits is shaped by such discordance (Mendes *et al*. 2018). This could in principle be accommodated in future extensions to the method by relaxing the number of topologies accepted as scaffolding trees, or by extending the model to accommodate such discordance.

Despite the potential utility in phylogenetics, there may be cases where useful phylogenetically-informative characters cannot be extracted from geometric morphometric data. This may be the case when any of the concerns stated by Felsenstein (2002) cannot be overcome, or when geometrically-defined characters exhibit inconsistent or weak signal, such as was found by Smith and Hendricks (2013) when using a semi-landmark geometric method to capture morphological variation in *Conus* snails. In these cases, it may be necessary to resort to using traditional linear measurements and proportions, or qualitative characters. Finally, there are cases where fossils may simply present weak information due to shortcomings in geologic and taxonomic sampling. When this is occurs, it is unlikely that any greater certainty in their placement can be achieved except by adding data.

### Comparison to other approaches

The method that I describe here differs substantially from existing approaches to the phylogenetic placement of fossil taxa. Although it is most similar to the fossil placement method developed by Revell *et al*. (2015), it extends their approach in several important ways. For instance, my approach does not require that branch lengths be scaled to time, simplifying the estimation procedure. In addition, the implementation here allows for the estimation of long extinct fossil taxa. Finally, the adaptation of Berger and Stamatakis’ approach to filtering character matrices can improve upon the accuracy achieved from existing methods. The method described here also differs from recent ’total-evidence’ methods that seek to simultaneously estimate both extinct and extant relationships. Although total-evidence methods are useful tools in the phylogenetic canon, splitting the estimation process into stages may be beneficial in certain datasets, and better suited to certain questions. For instance, the approach here may be used to generate priors for the placement of fossil calibrations in node dating. A new method has been developed that accommodates uncertainty in the phylogenetic placement of node calibrations in Bayesian molecular dating (Guindon 2018), which could, in principle, be combined with my fossil placement approach, by using posterior support of fossil calibrations as the prior probabilities in the dating analysis.

It is also worth noting that the method that I describe here would be straightforward to implement in existing phylogenetics packages, such as RevBayes, and adapted to a total-evidence framework. Although RevBayes does not feature a native implementation of the model that I describe, including the data-filtering approach, adapting the present procedure to this framework may be useful in addressing certain biological questions. This may include an exploration of the feasibility of incorporating continuous data into total-evidence morphological clock analyses (Zhang *et al*. 2015).

Moving forward, it will be important to explore the behavior of this method when applied to morphometric data collected under a variety of approaches and sampling schemes. The success of the weight calibrations on the simulated and empirical datasets suggests the possibility of applying the method to very large morphometric datasets by providing a means to filter through the noise that may occur when sampling densely across taxa and organs. Such a framework would facilitate the development of a more data-centric approach to morphological phylogenetics that reduces common sources of bias in morphological datasets by filtering data matrices statistically rather than through subjective judgement. This would encourage an exploration of conflict and concordance in signal through quantitative data analysis rather than by attempting to filter during the stage of data collection. They provide a means to move forward, and should encourage the development and exploration of more data sources to determine the most effective means to inform phylogeny using morphometry. One major gap in the approach presented here concerns the assumption that all continuous traits under analysis evolve under a shared rate. In the empirical analyses performed above, I rescaled the traits at each site so that the variance is set to be equal. However, moving forward, it will be important to explore model extensions that accommodate rate heterogeneity across characters. This has been done in continuous characters to positive effect by Schraiber *et al*. (2013) using a Gamma site-rate model, and adapting this or alternative approaches to modeling rate heterogeneity (Huelsenbeck and Suchard 2007) will be a key priority in extending the method described here moving forward.

## Conclusions

The method described here provides a new means for biologists to reliably and confidently place fossils in the tree of life. Although the simulated and empirical analyses show several imperfections and a need for further refinement of these methods, the overall accuracy and conservative assessment of uncertainty displayed in the examples appear encouraging. As molecular phylogenetics advances in its use of genomic data to answer fundamental questions across the tree of life, it will be important for morphological phylogenetics and paleontology to keep pace. Analysis of morphometric data using the approach shown here will help to improve issues surrounding subjectivity in character collection, and will help morphological datasets to scale better in the genomic era. New advances in the collection of morphometric data, combined with refinements to the approach developed here will better equip morphology to speak to major outstanding questions across the tree of life.

## Supplementary figures

**Figure S1.**
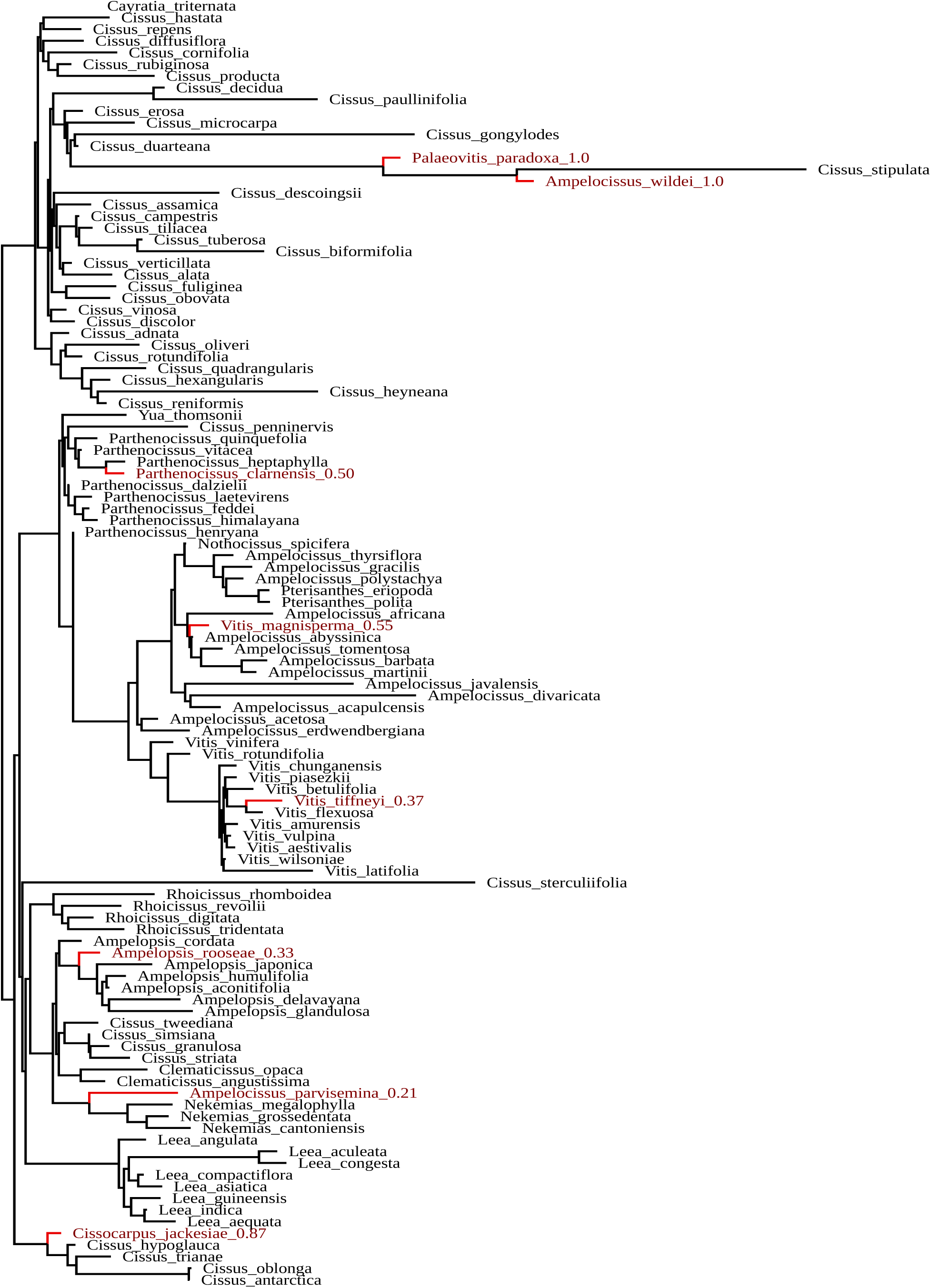
Vitaceae fossil placements inferred under the exponential branch length prior. Fossil taxa and branches are highlighted in red. Values following fossil tip labels indicate posterior support for placement. Topology is summarized from the posterior using the set of maximally credible clades (MCC). The placements are depicted only in the subclade containing all 6 fossils.

## Acknowledgements

I thank James Saulsbury, Gregory Stull, Joseph Walker, and Stephen Smith for helpful comments that improved the manuscript. I also greatly thank Iju Chen and Steven Manchester for providing the Vitaceae measurement data, and Gregory Stull for coordinating and helping to prepare the data for analysis.***Supplementary Data*** All supplemental material, including scripts, newick files, morphological and molecular alignments used in this study are available as a GitHub repository (https://github.com/carolinetomo/fossil_placement_tests).

